# An *in vitro* grafting method to quantify mechanical forces of adhering tissues

**DOI:** 10.1101/2020.08.21.260562

**Authors:** Yaichi Kawakatsu, Yu Sawai, Ken-ichi Kurotani, Katsuhiro Shiratake, Michitaka Notaguchi

## Abstract

Grafting is an indispensable agricultural technology for propagating useful tree varieties and obtaining beneficial traits of two varieties/species—as stock and scion—at the same time. Recent studies of molecular events during grafting have revealed dynamic physiological and transcriptomic changes. Strategies focused on specific grafting steps are needed to further associate each physiological and molecular event with those steps. In this study, we developed a method to investigate the tissue adhesion event, an early grafting step, by improving an artificial *in vitro* grafting system in which two pieces of 1.5-mm thick *Nicotiana benthamiana* cut stem sections were combined and cultured on medium. We prepared a silicone sheet containing five special cutouts for adhesion of cut stem slices. We quantitatively measured the adhesive force at these grafting interfaces using a force gauge and found that graft adhesion started 2 days after grafting, with the adhesive force gradually increasing over time. After confirming the positive effect of auxin on grafting by this method, we tested the effect of cellulase treatment and observed significant enhancement of graft tissue adhesion. Compared with the addition of auxin or cellulase individually, the adhesive force was stronger when both auxin and cellulase were added simultaneously. The *in vitro* grafting method developed in this study is thus useful for examining the process of graft adhesion.

## Introduction

Plant grafting has been used for thousands of years to improve crop traits and propagate fruit trees and vegetables. Wound healing at the graft junction allows two or more joined plants to grow as a single individual (Mudge et al. 2009; Melnyk 2016). Grafting confers various characteristics of roots (stocks), such as disease resistance and tolerance to adverse soil conditions, on shoots (scions) that also have desirable traits (Mudge et al. 2009). The degree of callus formation often observed on the grafted surface is believed to affect successful contact (Sass 1932). In the model plant Arabidopsis, the grafting process includes the following steps: necrotic layer development, graft callus growth, differentiation of new vascular tissue within the scion, and complete vascularization between the scion and the stock (Flaishman et al. 2008).

In regard to molecular mechanisms, studies have revealed the activity of enzymes, such as peroxidase and catalase, during grafting (Fernández-garcía et al. 2004; Pina and Errea 2008; Irisarri et al. 2015). Molecular genetic studies of grafting have been conducted in fruit trees, vegetables, and model plants (Cookson et al. 2014; Wang et al. 2014; Liu et al. 2015; Melnyk et al. 2015; Irisarri et al. 2015; Li et al. 2016; Matsuoka et al. 2016; Chen et al. 2017; Melnyk et al. 2016; Matsuoka et al. 2018; Wang et al. 2019; Xie et al. 2019; Kurotani et al. 2020; Notaguchi et al. 2020). Among the various findings, these studies have clearly shown that grafting is promoted by auxin action through the expression of genes related to auxin signaling. The phytohormone auxin is transported basipetally from synthetic tissues, such as leaves and buds, by polar transport and accumulates at the damaged site (Yin et al. 2012; Wang et al. 2014; Liu et al. 2015; Melnyk et al. 2015; Matsuoka et al. 2016; Chen et al. 2017; Melnyk et al. 2016; Kurotani et al. 2020; Notaguchi et al. 2020). Auxin promotes the formation of vascular strands when applied exogenously to undifferentiated tissue (Parkinson and Yeoman 1982; Wang et al. 2014). Recently, the graft adhesion capability of *Nicotiana benthamiana* (*Nb*) to phylogenetically distant plant species has been reported, and tomato fruits were produced on non-solanaceous plant families using a *Nb* stem as an interscion. This grafting ability is due to the *GH9B3* clade of β-1,4-glucanases secreted into the extracellular region (Notaguchi et al. 2020). Auxin and β-1,4-glucanases are thus good candidates for enhancing grafting techniques.

To measure the effect of the candidate compounds on graft promotion, the development of appropriate strategies to quantify their influence is desirable. *In vitro* micro grafting (IVG) is a simplified method for analyzing grafting traits. IVG involves the assembly of two explanted stem pieces and culturing on medium using a special mold or similar device to stabilize the positions of the tissues. Using this method, grafted seedlings are cultivated in a specific environment, and the physiological bond between stock and scion is examined (Parkinson and Yeoman 1982; Richardson et al. 1996; Ramanayake and Kovoor 1999; Dobránszki et al. 2000; Estrada-Luna et al. 2002; Raharjo and Litz 2005). IVG experiments have been conducted using a variety of plants, such as solanaceous species, apple, cacti, avocado, and prickly pear cactus, and the grafting efficiency has been evaluated by scoring success rates, measuring mechanical strengths, or observing the morphology of graft connections. IVG methods have also been used to examine the effects of phytohormones (Parkinson and Yeoman 1982; Dobránszki et al. 2000). In IVG experiments using solanaceous species, for example, two explanted internodes 7 mm long were inserted into a silicone tube and placed between two agar blocks. The addition of auxin to the agar medium resulted in the promotion of grafting (Parkinson and Yeoman 1982). In the described study, four IVG samples were tested per Petri dish. Thus, IVG methods are useful for examining the effect of chemical treatment on grafting in a small space without the use of entire plant individuals.

In the present study, we modified an IVG method and increased its throughput. We used *Nb* as a plant material because of its high grafting capability and the wide availability of various methodological and molecular data related to grafting in this species. To compare the effects of different grafting conditions, we quantitatively measured the extent of graft adhesion with a force gauge. Using this new method, we also investigated the effects of temperature and treatment with auxin and cellulases.

## Materials and Methods

### Plant materials

Sterilized seeds of *Nb* were grown on half-strength Murashige–Skoog (MS) gellan gum medium and cultivated under light at 27 °C for a week. The seedlings were then transferred to soil (1:1 mixture of vermiculite and Hanachan culture soil [Hanagokoro, Japan]) and grown for 3 to 4 weeks at 27 °C under continuous light conditions.

### Production of *in vitro* grafting sheets

Poly(dimethylsiloxane) (PDMS) sheets were created by mixing 30 g of SILPOT 184 W/C base (Dow Chemical, USA) with 1 g of catalyst in a 100-mL disposable cup (AS ONE, Japan). After depressurization in a desiccator for 20 min to remove air bubbles, the mixture was poured into a square Petri dish (100 × 100 × 15 mm; Simport Scientific, Canada), degassed in a desiccator for 20 min, and baked at 65 °C for 90 min. The resulting PDMS sheets (ca. 3-4 mm thickness) were cut into 2 × 8 cm portions. In the center of each sheet, five cutouts were created along the long axis at 1.3-cm intervals. To form each cutout, three adjacent, partially overlapping holes were punched into the sheet with a disposable biopsy punch (4.0 mm diameter; Kai Medical, Japan), which resulted in the creation of two pairs of protrusions (see also Figure 1A, B). The length of each cutout was approximately 9 mm.

**Figure 1.**
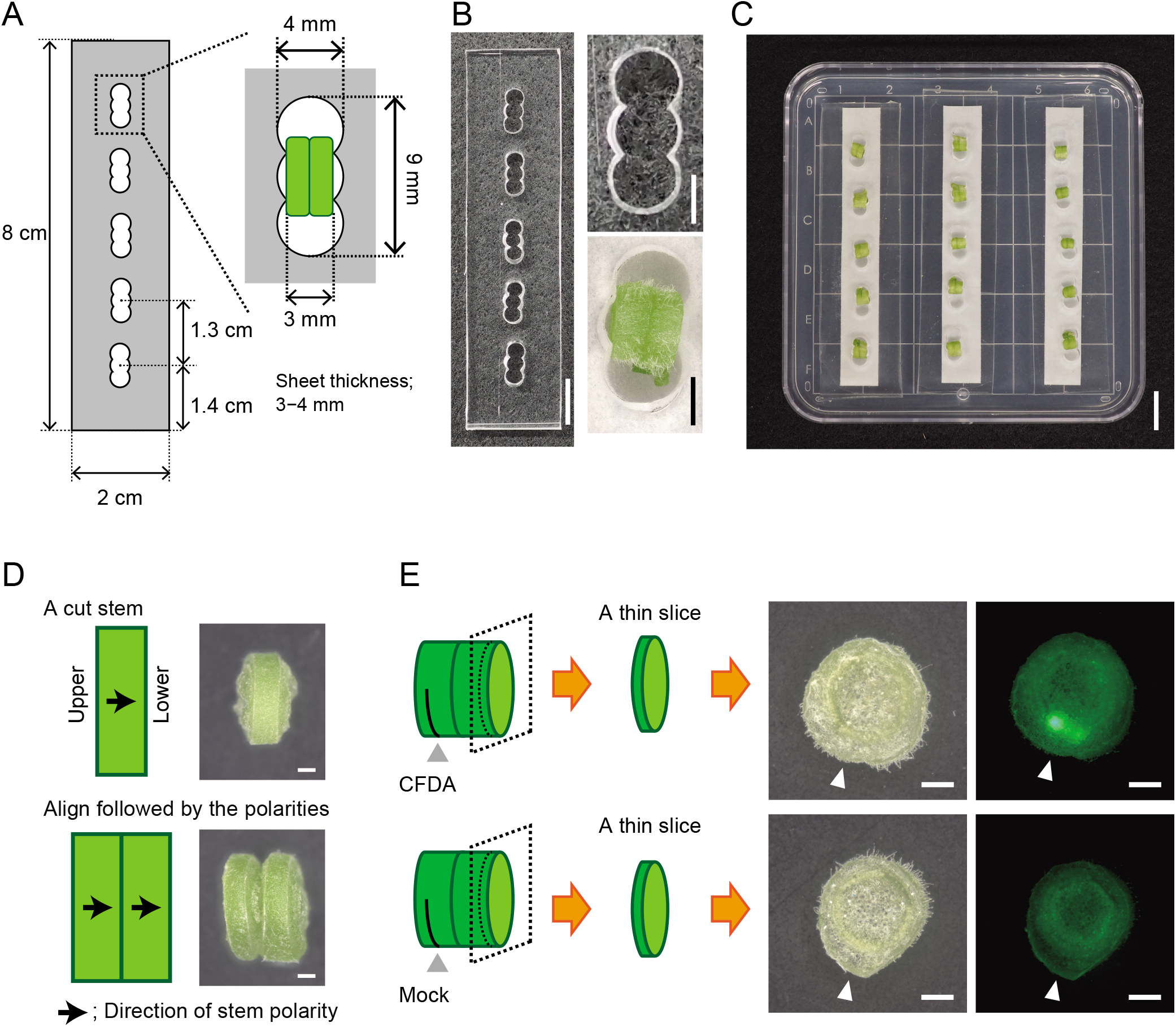
*In vitro* grafting (IVG) using a silicone sheet. (A) Design of an IVG grafting sheet. (B) Images of a sheet; an empty cutout is displayed in the upper right-hand corner, beneath which is shown a cutout holding a pair of IVG stem tissues. (C) Image of IVG culture in a dish. (D) Illustrations and photos taken 7 days after incubation of a piece of cut stem and a pair of IVG stems in which the original polarity, indicated by arrows, is preserved. (E) Tracer experiment for examining tissue connection 7 DAG. The results of treatment of one side of IVG tissues with CFDA or DMSO (mock) is shown. Thin slices sectioned from the other side of the tissues were examined for dye fluorescence. White arrowheads indicate the positions of applied CFDA and mock treatments. Scale bars: 1 cm (B and C), 1 mm (D and F).

### *In vitro* grafting

Autoclaved filter papers cut into 7.5 × 1 cm portions were placed on medium containing half-strength MS (pH 5.7), 1 % agar, and 300 ng mL^-1^ cefotaxime sodium salt (CSS) (Tokyo Chemical, Japan) prepared in a square Petri dish (100 × 100 × 15 mm; Simport Scientific). The *in vitro* grafting sheets were placed over the filter papers. Next, stem tissues were prepared as follows. Approximately 10–15cm lengths of *Nb* stems were excised from the pot-grown plants, sterilized with 70 % EtOH for 30 s, and rinsed three times with sterile water. After removal of both ends of the stem (ca. 0.5–1 cm) on a sterile filter paper, the stems (3–6 mm diameter) were cut into five to six 3-mm-wide pieces. Each stem tissue was then cut horizontally again to form two 1.5-mm-wide tissue slices. Two sliced tissue sections were placed against each other while preserving the original polarity of the donor plant and inserted together into one the cutouts on the *in vitro* grafting sheet. The dish containing the *in vitro* grafting sheets, usually three sheets or fewer, was covered and sealed with surgical tape and incubated at 27 °C under continuous light conditions. For testing the effects of auxin and cellulases, culture media were prepared by adding suitable amounts of 2,4-dichlorophenoxyacetic acid (Wako, Japan) and/or Onozuka R-10 cellulase mixture (Yakult Pharmaceutical, Japan) stock solutions dissolved in sterile water on the surfaces of the media, which was followed by drying under a clean bench for more than 30 min.

### Evaluation of IVG using tracer dyes

IVG stems were incubated for 7 days on a medium containing half-strength MS (pH 5.7), 2 % agar, and 300 ng mL^-1^ CSS. The symplasmic tracer 5(6)-carboxyfluorescein diacetate (CFDA; 50 mg mL^-1^ in dimethyl sulfoxide [DMSO]; Sigma, USA) was diluted with sterile water to 500 μg mL^-1^ to generate a working solution. One side of each IVG stem tissue was partly cut with a scalpel whose tip had been previously dipped in the CFDA working solution. After incubation of the IVG stem tissues under dark conditions at room temperature for 1 h, the non-cut side of each stem was sectioned for observation. The carboxyfluorescein fluorescence images were captured using an on-axis zoom microscope (Axio Zoom.V16, Zeiss, Germany) equipped with a digital camera (Axiocam 506 color, Zeiss). As a control, the same experiment was performed with 1 % aqueous DMSO without CFDA.

### Adhesive force measurements with a force gauge

As shown in Figure 2A–C, a looped string was attached with adhesive tape to the lid of a 35-mm Petri dish (Iwaki, Japan) prior to measurement. For easy displacement of samples from dish surfaces, a piece of adhesive tape (1 cm in diameter) was affixed to the center of each Petri dish. The Petri dish and its lid served as lower and upper parts, respectively, of the setup. Using instant glue (Aron Alpha; Toa Gosei, Japan), a set of IVG stem sections was then attached to the surface of the tape on the Petri dish and the lid. Approximately 30 s after glue attachment, the adhesive force of the IVG stem section was measured with a digital force gauge (ZTA-5N, Imada, Japan) set on a measuring stand (MX-500N, Imada). A looped string attached to the lid was connected to the force gauge, and the position of the Petri dish was fixed on the stage by hand. The force gauge was lifted up at a constant speed (5 mm s^-1^) until the set of IVG stem sections separated into two sections. Adhesive force was measured by subtracting the minimum value measured at the starting point. Stems that came apart on the *in vitro* grafting sheets during incubation were not measured. To test the correlation between graft area and adhesive force, photos of the graft interface of both upper and lower IVG stem tissues were taken after adhesive force measurements. The average value of the two stem areas was calculated as the adhesion surface area. A Mann–Whitney *U* test was used for comparisons of two data. Multiple comparisons of normally and non-normally distributed data were carried out using Tukey’s honestly different (HSD) and Steel–Dwass tests, respectively.

**Figure 2.**
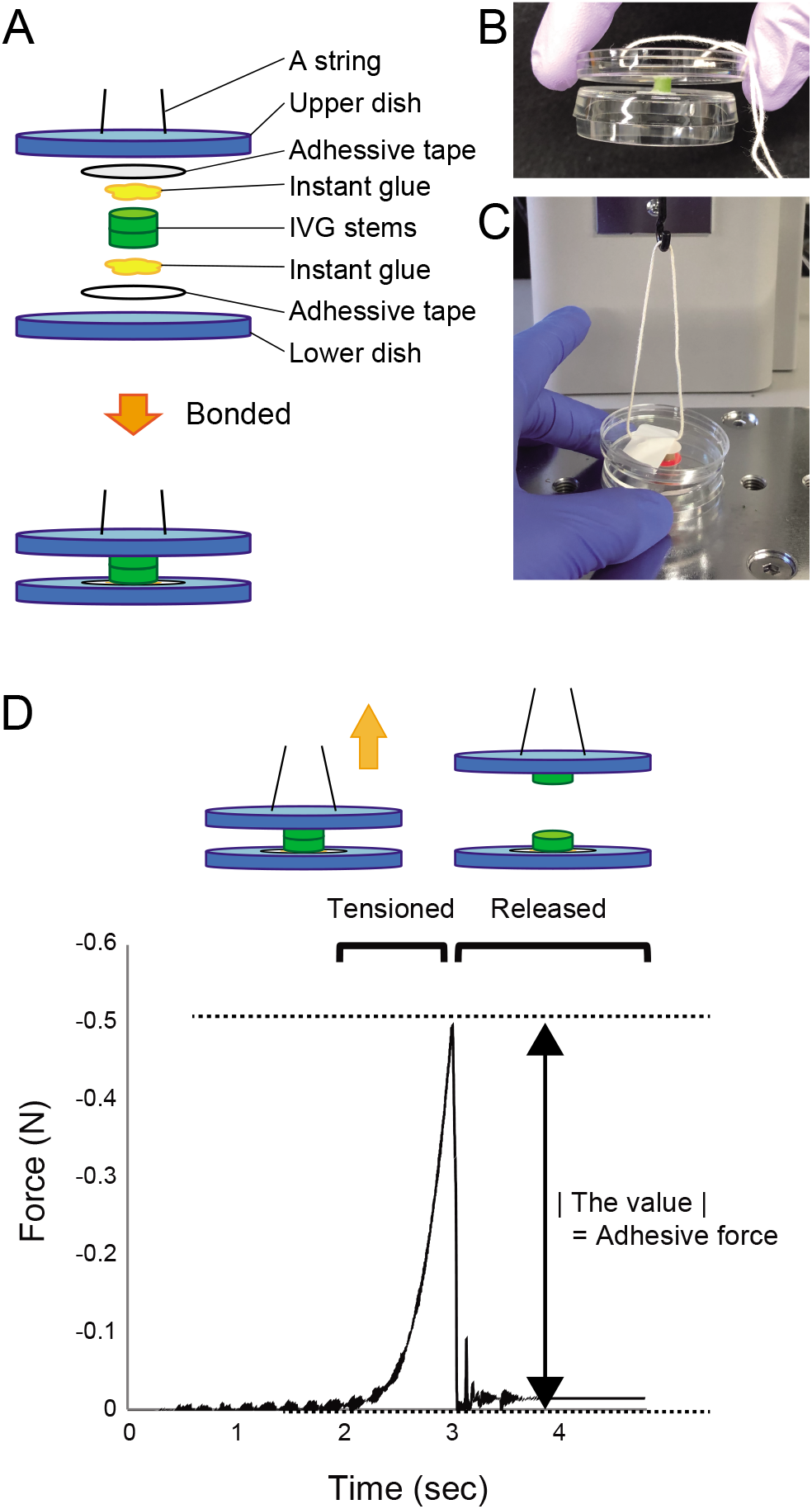
Measurement of the adhesive force of IVG stems using a force gauge. (A) Illustration of IVG sample preparation for force gauge measurement. (B) Image of IVG stems attached to a 35-mm dish as shown in A. (C) Image of attachment of *Nb* IVG stems to the force gauge. (D) Example of adhesive force measurement with the force gauge using an IVG sample at 7 DAG. The force value increased as the tension was increased until the *Nb* IVG stems separated (illustrated in upper panel). The absolute value of the difference between maximum and minimum values was defined as the adhesive force.

## Results and Discussion

### Development of the IVG method using a silicone sheet

IVG sheets made of PDMS silicone rubber were created to hold cut stem sections uniformly. Each 8 × 2 cm sheet contained five evenly spaced cutouts (one graft tissue per one cutout) (Figure 1A, B). Each of the approximately 4 × 9 mm cutouts had two pairs of elastic protrusions to help support the plant tissue (Figure 1A, B). Extra space remaining in the cutout after insertion of plant tissues allowed sample size variation to be accommodated to some extent. Because the cut stem tissues used for IVG gradually expanded outwards during incubation and pressed against the edges, the cutouts were positioned to not interfere with each other.

IVG was performed by combining two *Nb* stem slices, each with a thickness of approximately 1.5 mm (Figure 1A, B). The IVG samples were inserted into the cutouts on the silicone sheets, which were placed on a filter paper laid on incubation medium (Figure 1C). The *Nb* cut stem sections formed callus at both cut surfaces and also accomplish adhesion to each other 7 days after incubation (Figure 1D). Tissue adhesion was confirmed by symplasmic dye tracer experiments. When one side of an IVG stem tissue was treated with CFDA, fluorescence was detected on the other side (Figure 1E).

### Adhesive force measurements using a force gauge

The adhesive force of *Nb* IVG stem tissues was measured using a force gauge (see Materials and Methods for details). Both sides of IVG stem tissues were tightly fixed to the surfaces of plastic Petri dishes with instant adhesive glue (Figure 2A, B). A string already attached to the Petri dish lid was set on the hook of the force gauge, and the dish was manually positioned on the base (Figure 2C). The string was then pulled upward by the force gauge at a constant speed of 5 mm s^-1^. The change in the force value was recorded in real time (Figure 2D). The absolute value of the difference between the final value and that at the starting point was defined as the adhesive force of the IVG sample (Figure 2D). The accuracy of this method was also confirmed by testing the magnetic force. Repeated measurements using three different-strength magnets resulted in a significant change in the magnetic force (Supplementary Figure 1).

### Measurement of the adhesive force of IVG stems under different cultivation conditions

Using the above method, the adhesive strength of *Nb* IVG stems was measured 1 to 9 days after grafting (DAG). Adhesion between *Nb* IVG stem pieces was detected starting at 2 DAG (Figure 3A, Supplementary Table 1). The average adhesive force continued to increase through 9 DAG and did not reach a plateau within this period (Figure 3A, Supplementary Table 1). These results are consistent with a previous observation that the mechanical strength of IVG stems reached a maximum by 14 DAG (Parkinson and Yeoman 1982). Because high temperature often increases endogenous auxin synthesis and enhances graft efficiency (Gray et al. 1998; Turnbull et al. 2002; Tsutsui et al. 2020), we also examined the effect of temperature (22 °C vs. 27 °C) on *Nb* IVG. The adhesive force of *Nb* IVG stems was 0.25 ± 0.22 N (average ± standard deviation [SD], *n* = 23) at 22 °C and 0.26 ± 0.29 N (*n* = 26) at 27 °C (Figure 3B), which was not significantly different according to a Mann–Whitney *U* test *(P* = 0.49). This result appears to be reasonable, as major sites of auxin synthesis, such as the shoot apical region and leaves, are removed from tissue samples in IVG. We next examined the effect of exogenously applied auxin on IVG tissues. Compared with non-treated samples, adhesive force was significantly increased when *Nb* IVG stems were incubated on medium containing 0.5 μM auxin: 0.78 ± 0.39 N (*n* = 17) vs. 0.49 N ± 0.39 N (*n* = 19) in treated vs. untreated samples, respectively (Figure 3C). The promoting effects of auxin treatment in IVG have also been observed in previous grafting studies (Parkinson and Yeoman 1982).

**Figure 3.**
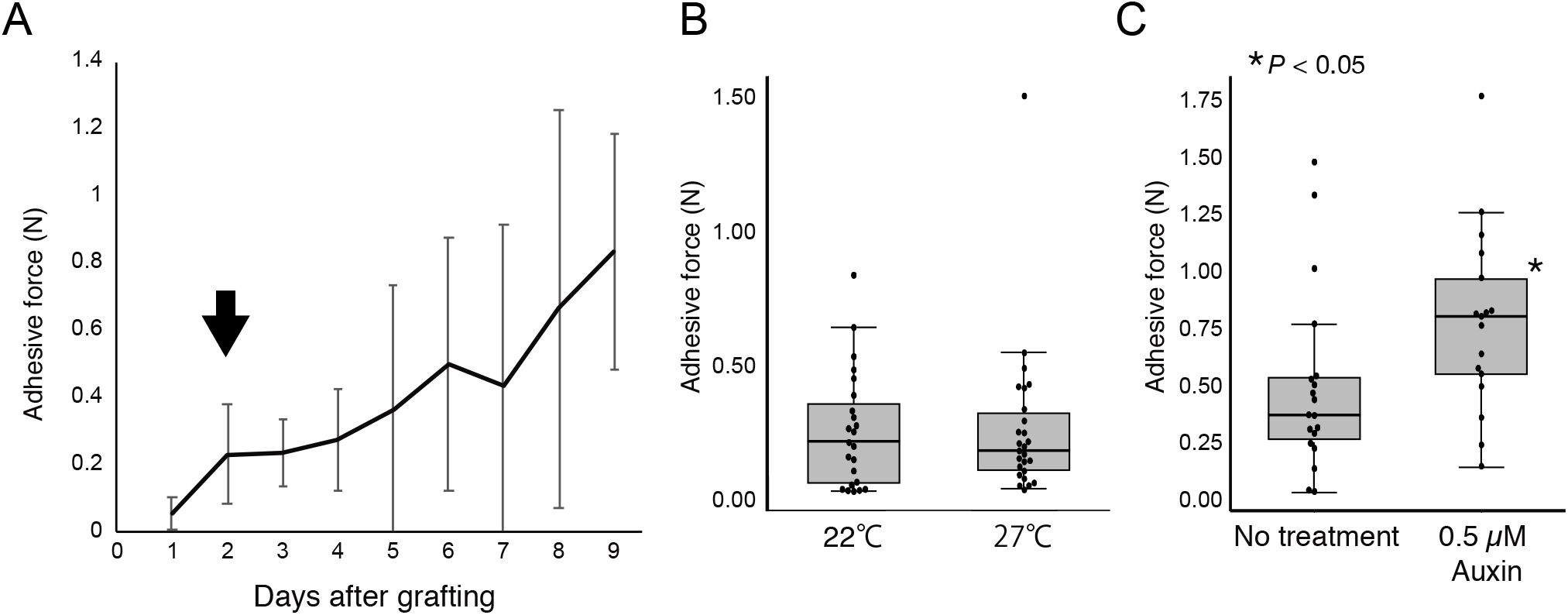
Tissue adhesion development in IVG. (A) Increase in adhesive force of *Nb* IVG stems 1 to 9 DAG. Error bars indicate SD. Adhesion was observed starting 2 DAG (arrow). (B) Adhesive force of *Nb* IVG stems incubated at 22 °C and 27 °C. (C) Adhesive force of *Nb* IVG stems incubated with or without auxin in the medium. An asterisk indicates statistical significance (*P*<0.05, Mann–Whitney *U* test).

### Relationship between adhesive force and grafting surface area

We next examined the relationship between adhesive force and grafting surface area, the latter calculated as the average of the areas of top and bottom stem surfaces attached to each other before force measurements (Figure 4A). In the case of both auxin-treated and untreated samples (same data as in Figure 3C), no correlation was identified between adhesive force and area (Figure 4B). When we divided the value of the adhesive force by the adhesive area (Figure 4C), we detected the same significant effect of auxin treatment shown in Figure 3C. Because tissue connection at the grafting junction is believed to take place in a region of the cambial tissues in the stem, a good fit between the cambial tissues of the stock and scion is crucial for grafting success (Mudge et al. 2009; Melnyk 2015). In our study, accordingly, the area of the graft interface did not exactly correspond to the area where the tissues were actually bonded. Because we were able to detect a significant effect of auxin treatment using the value of the adhesive force alone, we evaluated the extent of IVG adhesion using this value without considering area.

**Figure 4.**
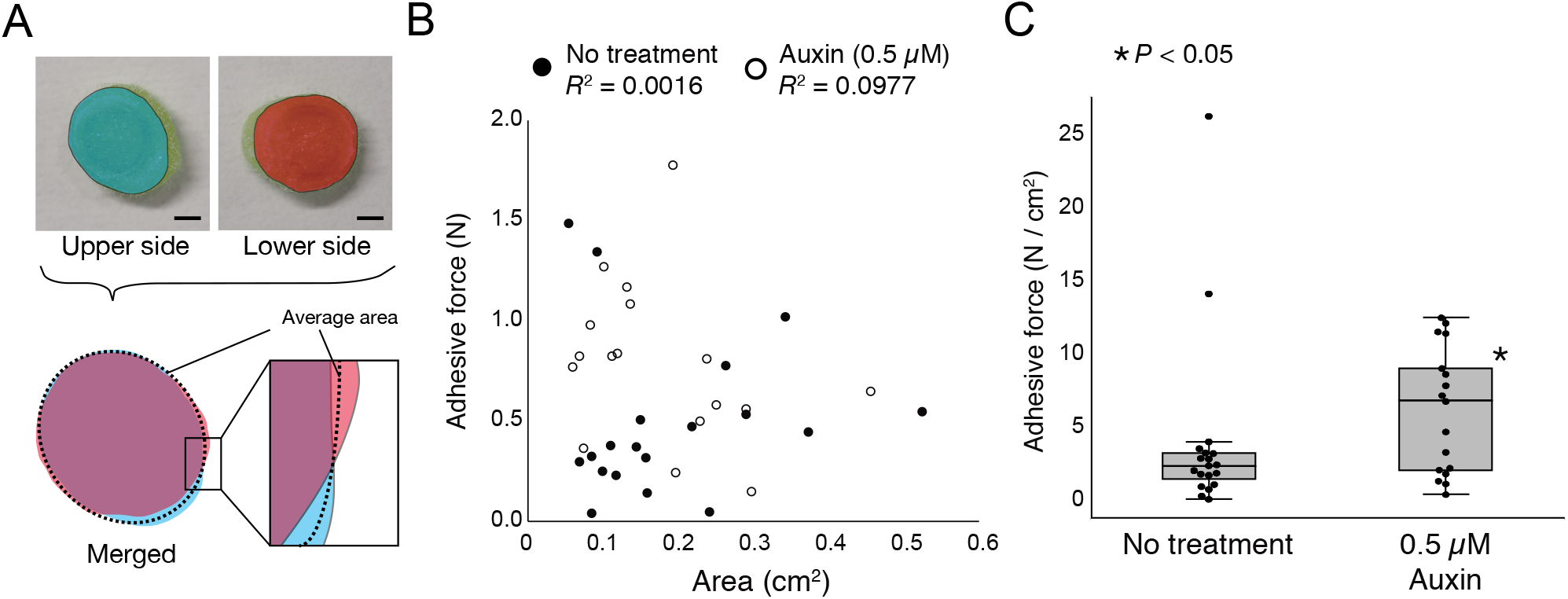
Relationship between graft interface area and adhesive force. (A) Measurement of the area of the grafted interface of *Nb* IVG stems. The graft area was calculated as the average of upper and lower tissue areas (dotted line). Scale bars, 5 mm. (B) Analysis of the correlation between the graft area and adhesive force of auxin-treated and untreated *Nb* IVG stems (the same stems represented in Figure 3C). In both cases, no correlation was observed between adhesive force and graft area. *R*^2^ is the coefficient of determination. (C) Box plot of adhesive force divided by each corresponding graft area (cm^2^) for the dataset in B. An asterisk indicates statistical significance (*P*<0.05, Mann–Whitney *U* test).

### Effect of cellulases on the adhesion of *Nb* IVG

We recently uncovered a key role of the *GH9B3* clade of β-1,4-glucanases in cell–cell adhesion at the graft junction (Notaguchi et al. 2020; Kurotani et al. 2020). β-1,4-glucanases probably target cellulose in cell walls, and the overexpression of the β-1,4-glucanase gene promotes graft efficiency in Arabidopsis (Notaguchi et al. 2020). In this study, we therefore tested whether exogenous application of cellulases facilitates graft adhesion using the *Nb* IVG system. We used a commercial, microorganism-produced cellulase mixture, henceforth referred to simply as cellulases. *Nb* IVG stems were incubated on medium containing cellulases at concentrations of 0.02 % and 0.2 %. Compared with non-treated samples, an increase in adhesive force was detected under both concentrations, with the stronger effects observed with 0.2 % cellulases (Figure 5A, Supplementary Table 2). Further effects were observed with 2.0 % cellulase application (Figure 5B). The measured adhesive force under normal medium and 2.0 % cellulase conditions was 0.24 ± 0.24 N (*n* = 23) and 0.82 ± 0.40 N (*n* = 27), respectively (Figure 5B). These results indicate that exogenous application of cellulases to tissues can promote grafting. As in the case of the auxin treatment shown in Figure 4, no correlation was found between adhesive force and adhesive area (Figure 5C). A significant effect due to cellulase treatment was also detected when the value of the adhesive force was divided by the adhesive area (Figure 5D).

**Figure 5.**
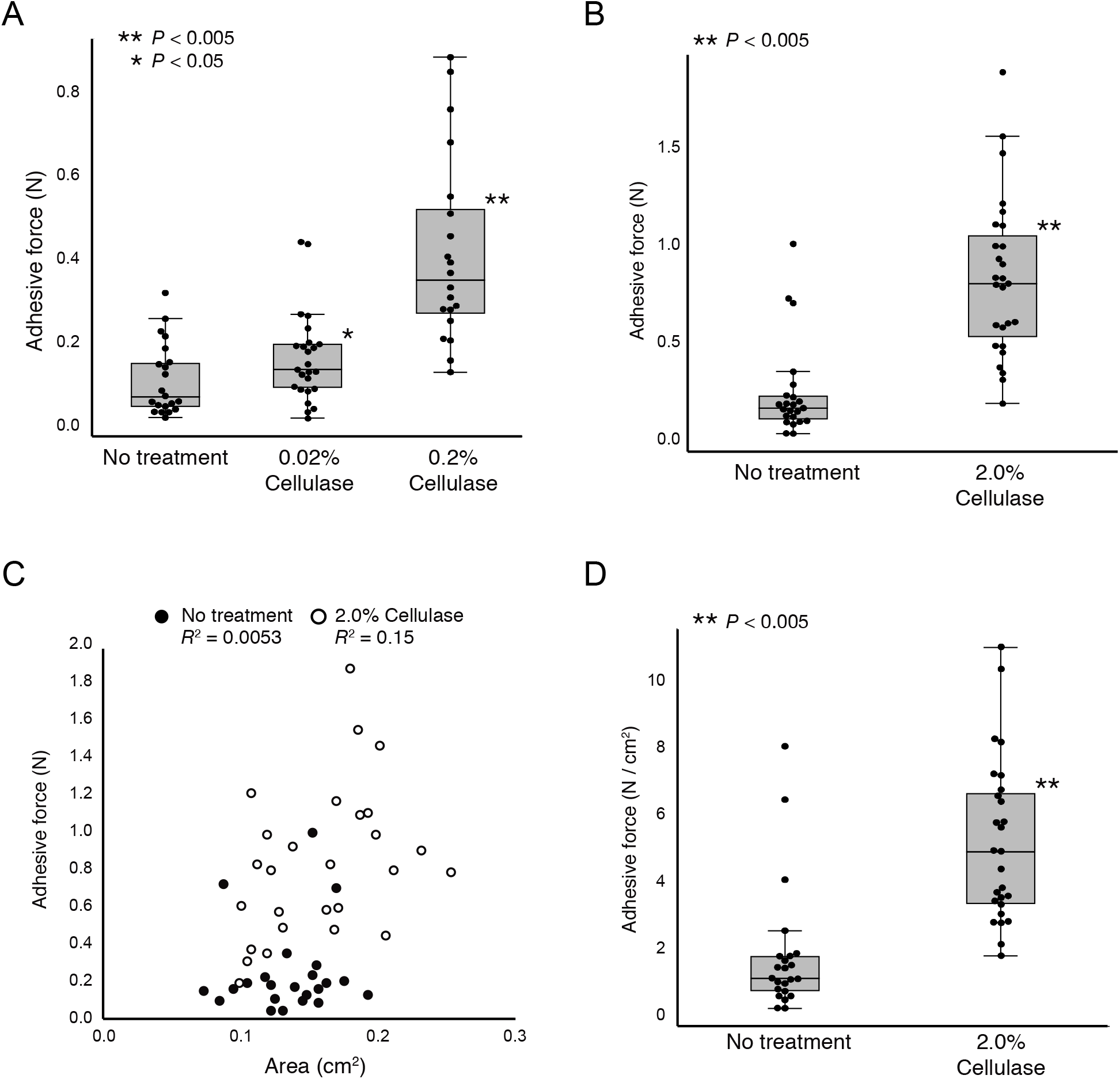
Effect of cellulase treatment on tissue adhesion in *Nb* IVG. (A, B) Adhesive force of *Nb* IVG stems incubated on cellulase-containing medium. (C) Analysis of the correlation between graft area and adhesive force of the *Nb* IVG stem datasets in B. In both cases, no correlation was observed between adhesive force and graft area. *R*^2^ is the coefficient of determination. (D) Box plot of adhesive force divided by each corresponding graft area (cm^2^) for the dataset in C. Asterisks indicate statistical significances (\**P* < 0.05, \*\**P* < 0.005, Mann–Whitney *U* test).

Finally, we tested whether the graft adhesion effects of auxin and cellulases were additive. We compared the results of no treatment, treatment with either 5 μM auxin or 0.2 % cellulase alone, and combined auxin–cellulase treatment. Compared with non-treated samples, an increase in adhesive force was observed in both auxin and cellulase individual treatments (Figure 6A, Supplementary Table 3). Combined treatment with auxin and cellulases resulted in a greater increase in adhesive force compared with either individual treatment (Figure 6A, Supplementary Table 3). Because the ranges of the generated data tended to vary, we plotted all the datasets on a logarithmic scale (Figure 6B). The data exhibited normal distributions; we therefore conducted a Tukey’s HSD test and confirmed the same tendency (Figure 6B). Overall, these results indicate that the promotion of grafting by auxin and cellulase takes place through two different mechanisms: auxin promotes cell division and callus formation at the graft junction, which enhances graft healing, whereas cellulases digest cellulose in cell walls and promote cell–cell adhesion at the graft interface. Auxin and cellulases can thus be applied together to achieve a synergic effect on graft enhancement.

**Figure 6.**
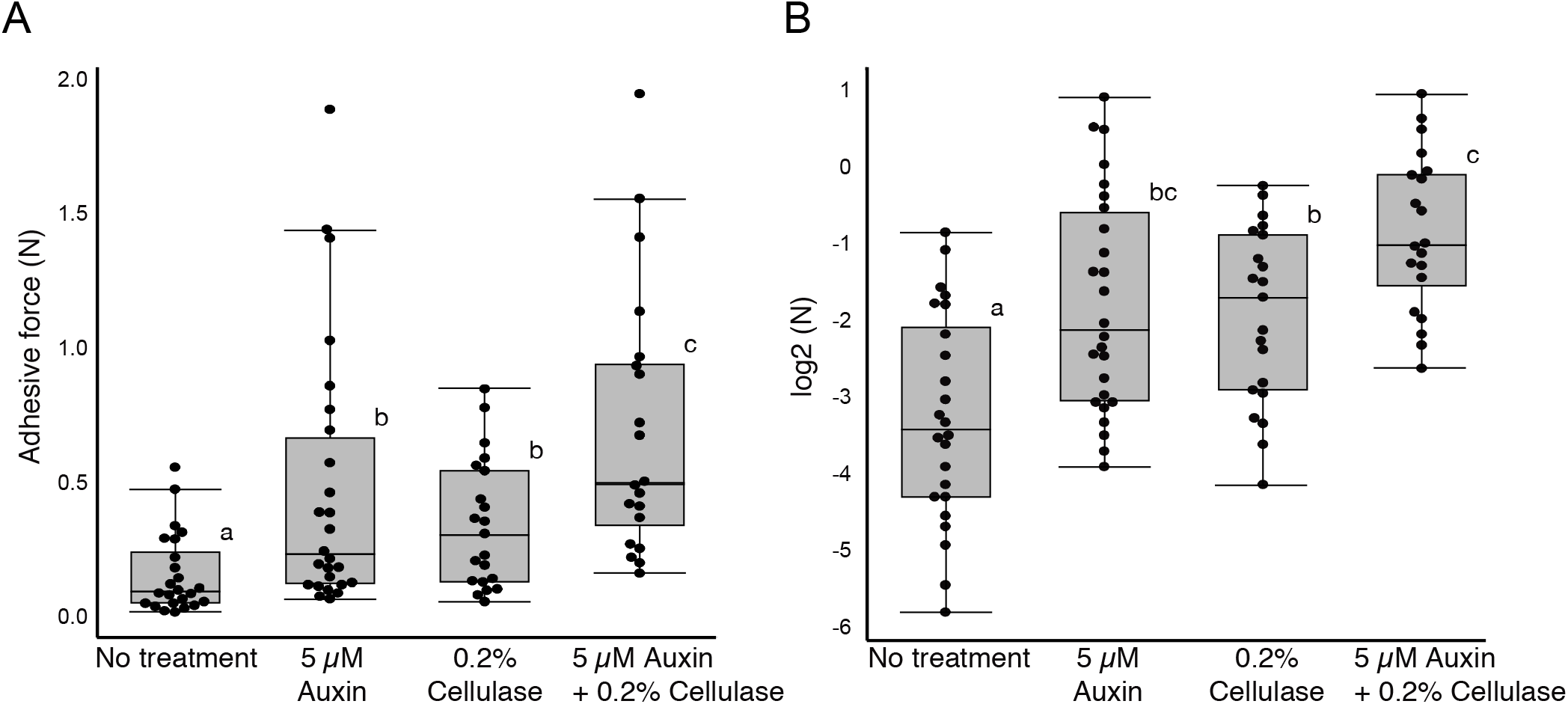
Additive effect of auxin and cellulase treatments on *Nb* IVG. (A) Adhesive force of *Nb* IVG stems incubated with either auxin or cellulase or both. The letters above boxes indicate statistical significance (*P* < 0.05, Steel–Dwass test). (B) Reevaluation of the data presented in A as plotted on a logarithmic scale. Normal distributions were observed. Promotion of grafting by auxin and cellulases was significant according to a Tukey’s HSD test (*P* < 0.05).

In this study, we developed a system for quantification of graft adhesion based on the IVG method. The modified IVG system has several advantages, such as high throughput, small space requirements and the ability to quantify adhesive force, which is critical for successful graft establishment. Using this system, we first showed that exogenous treatment with cellulases promotes graft adhesion. Given that numerous candidate genes related to grafting have been identified through recent transcriptome analyses (Cookson et al. 2014; Liu et al. 2015; Melnyk et al. 2015; Li et al. 2016; Chen et al. 2017; Wang et al. 2019; Xie et al. 2019; Kurotani et al. 2020; Notaguchi et al. 2020), methods for evaluating graft efficiency are required to test the roles of such genes. The IVG system can be applied to test the effect of genetic lines, environmental factors, and chemical treatments. The improvement of grafting techniques may increase opportunities for use of a variety of plant resources.

## Acknowledgments

We thank H. Makino, M. Matsumoto, A. Yagi, I. Yoshikawa, and M. Taniguchi for technical assistance. This work was supported by grants from the Japan Society for the Promotion of Science Grants-in-Aid for Scientific Research (18KT0040 and 19H05361 to M.N. and 18H03950 to K.S. and M.N.), the Cannon Foundation (R17-0070 to M.N.), and the Project of the NARO Bio-oriented Technology Research Advancement Institution (Research Program on Development of Innovative Technology 28001A and 28001AB to K.S. and M.N.).

## Author Contributions

M.N., K.S., and Y.K. conceived this study. Y.S. performed primary experiments. Y.K. designed and conducted the main experiments with advice from K.K. and M.N. Y.K. and M.N. wrote the paper.

## Abbreviations

CFDA: carboxyfluorescein diacetate
CSS: cefotaxime sodium salt
DMSO: dimethyl sulfoxide
HSD: honestly significant difference
IVG: *in vitro* grafting
MS: Murashige and Skoog
N: newton
PDMS: poly(dimethylsiloxane)
SD: standard deviation

## Supplementary Files

Supplementary Table 1. Adhesive force of *Nb* IVG stems at 1 to 9 DAGs.

Supplementary Table 2. Effect of cellulase treatment on adhesive force in *Nb* IVG.

Supplementary Table 3. Effect of auxin and cellulase treatments on adhesive force in *Nb* IVG.

Supplementary Table 4. Statistics of log2 values of the adhesive force in *Nb* IVG presented in Figure 6B.

Supplementary Figure 1. Measurement of magnetic adhesive force with a force gauge. (A) A representative waveform obtained by force measurement. The vertical axis represents adhesive force, and the horizontal axis represents time. Three consecutive states are shown: “tensioned,” “detached, but still reflected by the magnetic force,” and “fully released.” (B) Adhesive force measurement with three different-strength magnets. Measurements were performed three times for each magnet. The statistical significance of comparisons of each other data was detected by a Mann–Whitney *U* test (*P* < 0.005).

## References

Chen Z, Zhao J, Hu F, Qin, Y, Wang X, Hu G (2017) Transcriptome changes between compatible and incompatible graft combination of *Litchi chinensis* by digital gene expression profile. Sci Rep 7, 3954

Cookson SJ, Clemente Moreno MJ, Hevin C, Nyamba Mendome LZ, Delrot S, Magnin N, Trossat-Magnin C, Ollat N (2014) Heterografting with nonself rootstocks induces genes involved in stress responses at the graft interface when compared with autografted controls. J Exp Bot 65, 2473–2481

Dobránszki J, Magyar-Tábori K, Jámbor-Benczúr E, Lazányi J (2000) New in vitro micrografting method for apple by sticking. Int. J Hortic Sci 6, 79–83

Estrada-Luna AA, López-Peralta C, Cárdenas-Soriano E (2002) In vitro micrografting and the histology of graft union formation of selected species of prickly pear cactus (*Opuntia* spp.). Sci Hortic 92, 317–327

Fernández-García N, Carvajal M, Olmos E (2004) Graft union formation in tomato plants: peroxidase and catalase involvement. Ann Bot 93, 53–60

Flaishman MA, Loginovsky K, Golobowich S, Lev-Yadun S (2008) *Arabidopsis thaliana* as a model system for graft union development in homografts and heterografts. J Plant Growth Regul 27, 231

Gray WM, Östin A, Sandberg G, Romano CP, Estelle M (1998) High temperature promotes auxin-mediated hypocotyl elongation in Arabidopsis. Proc Natl Acad Sci USA 95, 7197–7202

Irisarri P, Binczycki P, Errea P, Martens HJ, Pina A (2015) Oxidative stress associated with rootstock–scion interactions in pear/quince combinations during early stages of graft development. J Plant Physiol 176, 25–35

Kurotani K, Wakatake T, Ichihashi Y, Okayasu K, Sawai Y, Ogawa S, Cui S, Suzuki T, Shirasu K, Notaguchi M (2020) Host-parasite tissue adhesion by a secreted type of β-1,4-glucanase in the parasitic plant *Phtheirospermum japonicum*. Commun Biol 3, 407

Li G, Ma J, Tan M, Mao J, An N, Sha G, Zhang D, Zhao C, Han M (2016) Transcriptome analysis reveals the effects of sugar metabolism and auxin and cytokinin signaling pathways on root growth and development of grafted apple. BMC Genomics 17, 150

Liu N, Yang J, Fu X, Zhang L, Tang K, Guy KM, Hu Z, Guo S, Xu Y, Zhang M (2015) Genome-wide identification and comparative analysis of grafting-responsive mRNA in watermelon grafted onto bottle gourd and squash rootstocks by high-throughput sequencing. Mol Genet Genomics 291, 621–633

Matsuoka K, Sugawara E, Aoki R, Takuma K, Terao-Morita M, Satoh S, Asahina M (2016) Differential cellular control by cotyledon-derived phytohormones involved in graft reunion of Arabidopsis hypocotyls. Plant Cell Physiol 57, 2620–2631

Matsuoka K, Yanagi R, Yumoto E, Yokota T, Yamane H, Satoh S, Asahina M (2018) RAP2.6L and jasmonic acid–responsive genes are expressed upon Arabidopsis hypocotyl grafting but are not needed for cell proliferation related to healing. Plant Mol Biol 96, 531–542

Melnyk CW (2016) Plant grafting: insights into tissue regeneration. Regeneration 4, 3–14

Melnyk CW, Schuster C, Leyser O, Meyerowitz EM (2015) A developmental framework for graft formation and vascular reconnection in *Arabidopsis thaliana*. Curr Biol 25, 1306–1318

Mudge K, Janick J, Scofield S, Goldschmidt EE (2009) A History of Grafting, in: Janick J. (eds) Horticultural Reviews. John Wiley & Sons Inc Hoboken, USA, pp 437–493

Notaguchi M, Kurotani K, Sato Y, Tabata R, Kawakatsu Y, Okayasu K, Sawai Y, Okada R, Asahina M, Ichihashi Y, Shirasu K, Suzuki T, Niwa M, Higashiyama T (2020) Cell-cell adhesion in plant grafting is facilitated by β-1,4-glucanases. Science 369, 698–702

Parkinson M, Yeoman MM (1982) Graft Formation in Cultured, Explanted Internodes. New Phytol 91, 711–719

Pina A, Errea P (2008) Influence of graft incompatibility on gene expression and enzymatic activity of UDP-glucose pyrophosphorylase. Plant Sci 174, 502–509

Raharjo SHT, Litz RE (2005) Micrografting and ex vitro grafting for somatic embryo rescue and plant recovery in avocado (*Persea americana*). Plant Cell Tissue Organ Cult 82, 1–9

Ramanayake SMSD, Kovoor A (1999) In vitro micrografting of cashew (*Anacardium occidentale* L.). J Hortic Sci Biotechnol 74, 265–268

Richardson FVM, tSaoir SMA, Harvey BMR (1996) A Study of the graft union in in vitro micrografted apple. Plant Growth Regul 20, 17–23

Sass JE (1932) Formation of callus knots on apple grafts as related to the histology of the graft union. Bot Gaz 94, 364–380

Tsutsui H, Yanagisawa N, Kawakatsu Y, Ikematsu S, Sawai Y, Tabata R, Arata H, Higashiyama T, Notaguchi M (2020) Micrografting device for testing systemic signaling in Arabidopsis. Plant J 103, 918–929

Turnbull CGN, Booker JP, Leyser HMO (2002) Micrografting techniques for testing long-distance signalling in Arabidopsis. Plant J 32, 255–26

Wang H, Zhou P, Zhu W, Wang F (2019) *De novo* comparative transcriptome analysis of genes differentially expressed in the scion of homografted and heterografted tomato seedlings. Sci Rep 9, 20240

Wang J, Jin Z, Yin H, Yan B, Ren ZZ, Xu J, Mu CJ, Zhang Y, Wang MQ, Liu H (2014) Auxin redistribution and shifts in PIN gene expression during Arabidopsis grafting. Russ. J. Plant Physiol 61, 688–696

Xie L, Dong C, Shang Q (2019) Gene co-expression network analysis reveals pathways associated with graft healing by asymmetric profiling in tomato. BMC Plant Biol 19, 373

Yin H, Yan B, Sun J, Jia P, Zhang Z, Yan X, Chai J, Ren Z, Zheng G, Liu H (2012) Graft-union development: a delicate process that involves cell–cell communication between scion and stock for local auxin accumulation. J Exp Bot 63, 4219–4232

