## Supplemental Figure 1 for "An *in vitro* grafting method to quantify mechanical forces of adhering tissues"

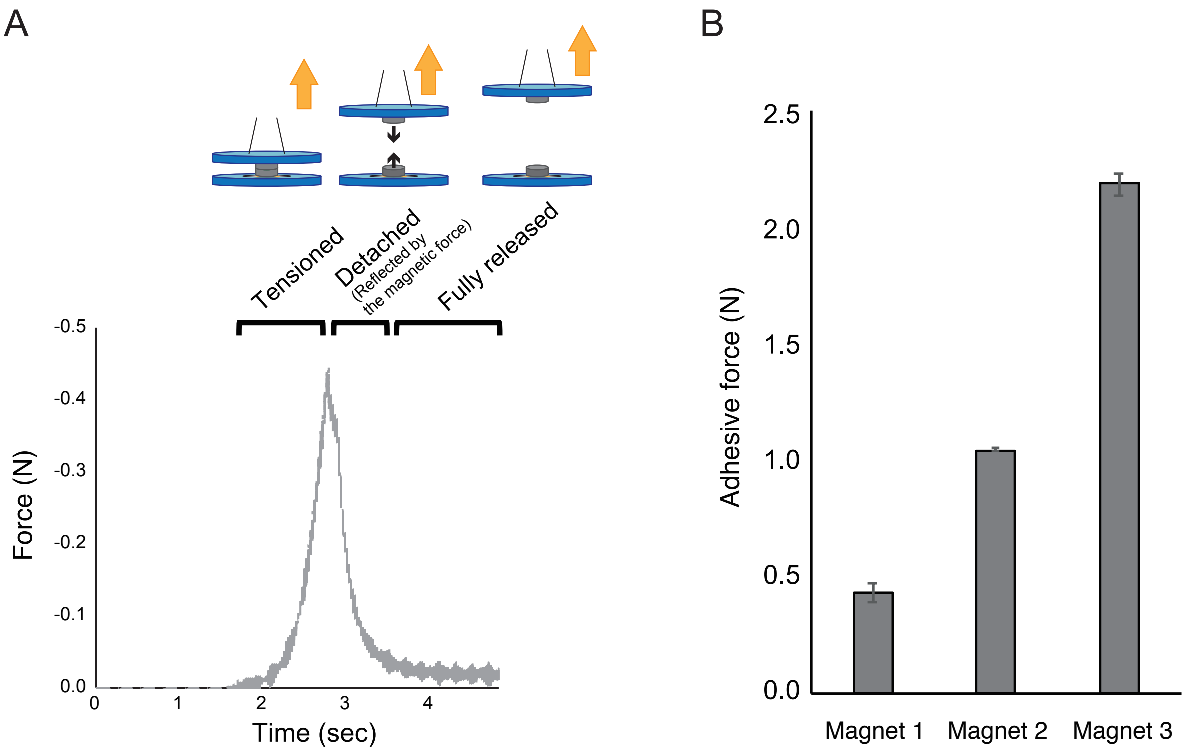


Supplementary Figure 1. Measurement of magnetic adhesive force with a force gauge. (A) A representative waveform obtained by force measurement. The vertical axis represents adhesive force, and the horizontal axis represents time. Three consecutive states are shown: "tensioned," "detached, but still reflected by the magnetic force," and "fully released." (B) Adhesive force measurement with three different-strength magnets. Measurements were performed three times for each magnet. The statistical significance of comparisons of each other data was detected by a Mann–Whitney *U* test ($P < 0.005$).
